# lociPARSE: a locality-aware invariant point attention model for scoring RNA 3D structures

**DOI:** 10.1101/2023.11.04.565599

**Authors:** Sumit Tarafder, Debswapna Bhattacharya

## Abstract

A scoring function that can reliably assess the accuracy of a 3D RNA structural model in the absence of experimental structure is not only important for model evaluation and selection but also useful for scoring-guided conformational sampling. However, high-fidelity RNA scoring has proven to be difficult using conventional knowledge-based statistical potentials and currently-available machine learning-based approaches. Here we present lociPARSE, a locality-aware invariant point attention architecture for scoring RNA 3D structures. Unlike existing machine learning methods that estimate superposition-based root mean square deviation (RMSD), lociPARSE estimates Local Distance Difference Test (lDDT) scores capturing the accuracy of each nucleotide and its surrounding local atomic environment in a superposition-free manner, before aggregating information to predict global structural accuracy. Tested on multiple datasets including CASP15, lociPARSE significantly outperforms existing statistical potentials (rsRNASP, cgRNASP, DFIRE-RNA, and RASP) and machine learning methods (ARES and RNA3DCNN) across complementary assessment metrics. lociPARSE is freely available at https://github.com/Bhattacharya-Lab/lociPARSE.

## 1 Introduction

Computational prediction of RNA 3-dimensional structures from nucleotide sequence has garnered considerable research effort over the past decade [1–10], and deep learning-enabled RNA 3D modeling has gained significant attention in the recent past [11–15]. To facilitate the practical applicability of predicted 3D models, it is critical to have a scoring function that can reliably assess their global topology and local quality in the absence of experimental structures [16–18]. Moreover, the ability of a scoring function to distinguish accurate 3D models of previously unseen RNAs from misfolded alternatives plays an important role in guiding conformation sampling towards the native state [19].

Existing methods for scoring RNA structures roughly belong to two categories: knowledge-based statistical potentials and supervised machine learning. Various knowledge-based statistical potentials have been developed, both at all-atom and coarse-grained levels [20–24], using different simulated reference states [25–30]. However, reliably distinguishing accurate structural models of RNA from less accurate ones has proven to be difficult because the characteristics of energetically favorable RNA structures are not sufficiently well understood and thus the reference states may deviate largely from the ideal one. Machine learning-based methods [17, 31] aim to overcome such limitations by learning to predict the accuracy of an RNA structural model through supervised learning. Indeed, machine learning-based RNA scoring functions, trained to estimate the unfitness score either at the nucleotide level or at the structural level by learning to predict the root mean square deviation (RMSD) from the unknown true structure, have been shown to be effective in RNA-Puzzles blind structure prediction challenges [5].

Despite the effectiveness, the existing machine learning methods do not consider some key factors that can significantly improve the sensitivity of RNA scoring functions. First, global superposition-dependent RMSD metric is not length normalized, is affected by superposition, is dominated by outliers in poorly modeled structural regions, and does not take into account the accuracy of the local atomic environment. RNA is a flexible molecule in which irregular loops may affect RMSD measures and global superposition may not be optimal, leading to scoring anomalies. Yet, virtually all existing machine learning-based RNA scoring functions use RMSD as the ground truth during supervised training. Second, similar to other macromolecules, RNA structures have no natural canonical orientation i.e., the same RNA structure can be rotated in space without affecting its biological function, thus allowing a structure to be represented in any orientation. As such, machine learning methods that are not invariant to global Euclidean transformations such as rotation must account for this aspect of variation by tweaking model architecture and/or parameters, which may affect their expressiveness and generalizability. Third, in consideration of RNA as a flexible molecule in which interplay between various local structural motifs defines the global topology, an effective scoring function should not be strongly influenced by the relative motions between the tertiary motifs. That is, the effects of relative movement between the motifs should not lead to artificially unfavorable scores.

Using the Local Distance Difference Test (lDDT) [32] as the ground truth during supervised training is an attractive alternative to the popular RMSD metric. lDDT compares distances between atoms that are nearby (within 15 Å) in the experimental structure to the distances between those atoms in the predicted structure and offers several advantages over RMSD. First, being superposition-free and based on rotation-invariant properties of a structure, lDDT naturally preserves invariance with respect to the global Euclidean transformations of the input RNA structure such as global rotations and translations. Second, lDDT measures the accuracy of the local environment of the model in atomic detail, without being affected by superposition or dominated by outliers in poorly modeled structural regions. Third, lDDT exhibits robustness to movements between tertiary structural units such as domains in proteins that can generalize to RNA tertiary motifs, provided a way can be found that ensures rigid motion between a set of local structural units is invariant under global Euclidean transformations on the said units. A solution to this problem comes from Invariant Point Attention (IPA) proposed in AlphaFold2 as part of the structural module [33]. IPA is a form of attention that acts on a set of 3D point clouds and is invariant under global Euclidean transformations on said points, where 3D point clouds are represented using local frames.

How can we capture the aforementioned benefits of lDDT in a neural network architecture for RNA scoring, while maintaining invariance under global Euclidean transformations? Here, we provide such a solution by developing a new attention-based architecture, called lociPARSE (<u>loc</u>ality-aware <u>i</u>nvariant <u>P</u>oint <u>A</u>ttentionbased <u>R</u>NA <u>S</u>cor<u>E</u>r), for scoring RNA 3D structures. Different from previous supervised learning approaches that estimate the RMSD metric, our method estimates local nucleotide-wise lDDT scores that are then aggregated over all nucleotides to predict global structural accuracy. Inspired by AlphaFold2, we define nucleotide-wise frames parameterized by rotation matrices and translation vectors operating on predefined RNA conformation at the local level. To model the local atomic environment of each nucleotide as captured by lDDT, the IPA implementation used in the original AlphaFold2 has been modified to incorporate locality information derived from the RNA atomic coordinates. By so doing, we are able to effectively capture the accuracy of each nucleotide while considering the effect of its local atomic environment. It is worth mentioning here that, although an RNA 3D structure prediction method that uses AlphaFold2-inspired IPA architecture can self-estimate the quality of its own predicted structure given an input RNA sequence, our method is the first general-purpose RNA scoring method that is capable of estimating the quality of any input RNA 3D structure using our modified implementation of AlphaFold2’s IPA architecture.

Our method significantly outperforms traditional knowledge-based statistical potentials as well as state-of-the-art machine learning-based RNA scoring functions such as ARES [31] on multiple independent test datasets including CASP15 blind test targets across a wide range of performance measures. In particular, lociPARSE exhibits superior ability to reproduce the ground truth lDDT scores both at the global and local levels, rank predictions for a given target with high fidelity, recognize the best predictions consistently, and better discriminate between ‘good’ and ‘bad’ predictions. An open-source software implementation of lociPARSE, licensed under the GNU General Public License v3, is freely available at https://github.com/Bhattacharya-Lab/lociPARSE.

## 2 Results

### 2.1 lociPARSE: locality-aware invariant point attention for RNA scoring

An overview of our method, lociPARSE, is illustrated in **Figure 1**. The core component of our architecture, outlined in **Figure 1b**, is an invariant point attention (IPA) module which utilizes the geometry of the input RNA 3D structure to revise the nucleotide and pair features. This component is similar to the AlphaFold2’s IPA formulation used in the structure module, but modified herein to incorporate locality information derived from the RNA atomic coordinates. To do this, we introduce locality-aware geometry and edge-biased attention (Section 4.2.2) to convert nucleotide pair features to edge adjacencies by considering a set k-nearest neighbor nucleotides to capture the local atomic environment of each nucleotide based on the Euclidean distances of the C4^*′*^ - C4^*′*^ atoms between nucleotide pairs. In our setting, we define local nucleotide frames (Section 4.2.1) from the Cartesian coordinates of C4^*′*^, P, and glycosidic N atoms. The IPA partitions the nucleotide query and value features into 3D vectors and transforms them from the target nucleotide’s local frame into a global reference frame before computing both attention weights and the output of the attention mechanism. Further, we augment nucleotide-nucleotide atomic distances between all pairs of 3 atoms P, C4^*′*^ and N, encoded with Gaussian radial basis functions as pair features (Section 4.1), and make further use of the pair features to bias attention weights and update scalar features. Our network architecture consists of 4 IPA layers, with the IPA hyperparameters (*N*_*heads*_, c, *N*_query points_, *N*_value points_) set to (4, 128, 8, 4) and we use 20 nearest neighbors for the locality computation, determined through ablation experiments using an independent validation set (Section 2.7). The output of the attention layer is invariant to the global Euclidean transformations such as global rotations and translations of the input RNA. Finally, a linear layer followed by a two-layer fully connected network is used to estimate the predicted nucleotide-wise lDDT (pNuL) scores. These nucleotide-level pNuL predictions are then aggregated over all nucleotides by taking an average to estimate the predicted molecular-level lDDT (pMoL), enabling our method to estimate both local and global qualities of the input RNA 3D structure.

**Figure 1:**
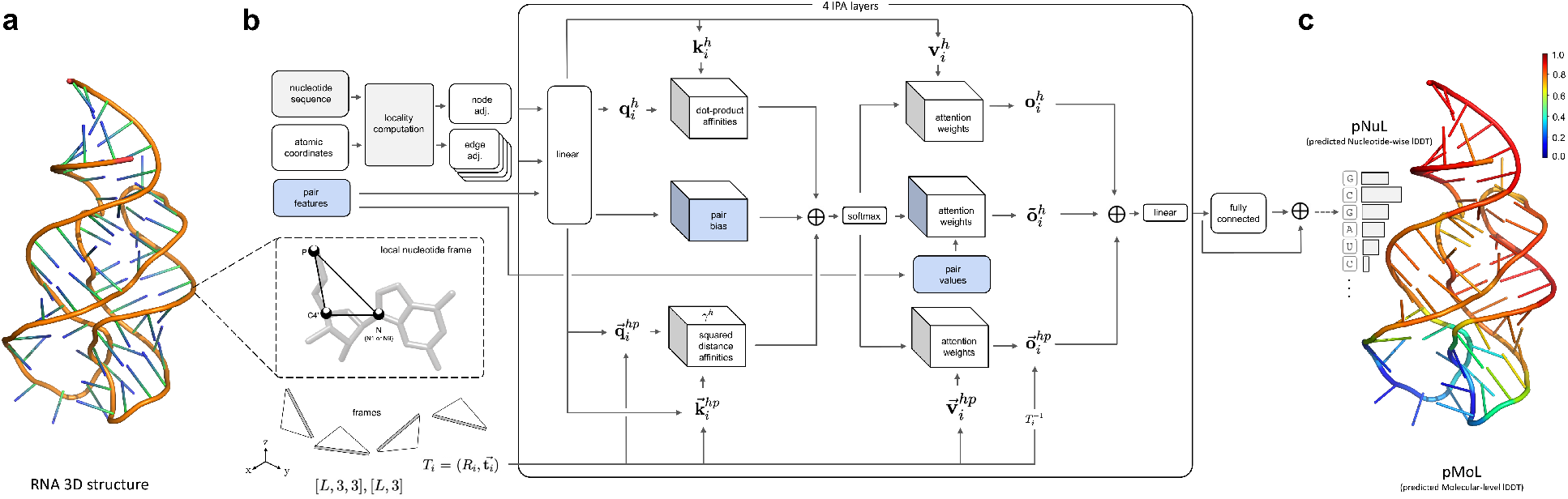
Overview of lociPARSE. Given a 3D RNA structure, we estimate the local nucleotide-wise lDDT scores that are then aggregated over all nucleotides to predict global structural accuracy. (a) An input RNA 3D structure. (b) The architecture of our locality-aware invariant point attention (IPA) module to capture the accuracy of each nucleotide and the effect of its local atomic environment. (c) By aggregating information at the level of nucleotide, we output predicted nucleotide-wise lDDT (pNuL) scores before making a prediction at the level of the entire RNA structure to output predicted molecular-level lDDT (pMoL).

### 2.2 Experimental setup and performance evaluation

For training and performance evaluation, we use existing and publicly available benchmark datasets. Our training dataset contains 1, 399 RNA targets collected from the training set used in the recent RNA 3D structure prediction method trRosettaRNA [12]. We extracted the sequences from the 1, 399 experimental structures and generated a total of 51, 763 structural models using a combination of different RNA 3D structure prediction tools including recent deep learning-enabled RNA structure prediction methods [11–15], physics-based RNA folding [34], and structure perturbation using PyRosetta [35] (Section 4.3). Details of our training procedure are provided in Section 4.4. Our test data includes 30 independent RNAs, also collected from trRosettaRNA following the train and test splits of the original work. We generated 3D structural models for each of these 30 RNAs using the deep learning-enabled RNA structure prediction methods [11–15]. We also use 12 RNA targets from CASP15 as an additional independent test set containing targets cleared for public access as of December 20, 2022, where the corresponding 3D structural models are collected directly from the CASP15 website https://predictioncenter.org/casp15/ based on the blind predictions submitted by various participating groups in CASP15 RNA 3D structure prediction challenge. We compare our method lociPARSE with traditional knowledge-based statistical potentials including rsRNASP [20], cgRNASP [21], RASP [22] and DFIRE-RNA [23] as well as state-of-the-art machine learning-based RNA scoring functions RNA3DCNN [17] and ARES [31]. To assess the accuracy of different aspects of quality estimation, we use a wide-range of performance measures including the ability to reproduce the ground truth lDDT and all-atom RMSD scores both at the global and local levels, rank predictions for a given target, recognize the best predictions, and discriminate between ‘good’ and ‘bad’ predictions. Our performance assessment can be broadly grouped into two categories: global level assessment and per-target average assessment. The global level assessment puts together the predicted scores of all the structural models across all targets during performance evaluation. In contrast, the per-target average assessment evaluates the predictions for each target’s structures separately against their corresponding ground truths and then averages the results over all the targets. Since the global level assessment puts all the targets together to evaluate, it is important to length-normalize the estimated scores for the methods that predict RMSD unfitness scores. For the same reason, we length-normalize the ground truth all-atom RMSD metric during scoring performance evaluation. For the length normalization of RMSD in the range of (0-1], we use the formulation of US-align [36] as follows:

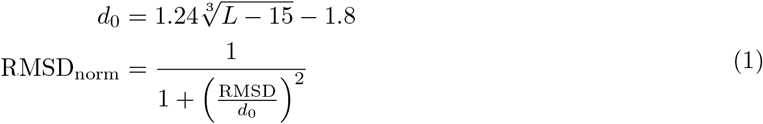

Except for lociPARSE, all the other six competing methods evaluated in this work estimate some form of structural unfitness score with a lower value representing better structural quality. ARES and RNA3DCNN estimate RMSD, whereas the four other knowledge-based statistical potentials estimate the potential energy of an RNA structural model. To ensure that the estimated scores from all competing methods are comparable, we length-normalize the predicted RMSD of ARES and RNA3DCNN using Equation 1 that maps the predictions between (0-1] with higher values representing better structural quality. For the knowledge-based statistical potentials, we use min-max normalization to map the estimated potential energy to a normalized score s between (0-1] and use (1− *s*) to re-calibrate the score such that a higher value represents better structural quality. We also length-normalize the ground truth RMSD between (0-1] using Equation 1. Our method lociPARSE estimates the lDDT score between (0-1] where higher values represent better structural quality. Meanwhile, the ground truth lDDT is between (0-1] by definition. Details of competing methods and evaluation metrics can be found in Section 4.5.

### 2.3 Performance on 30 independent RNA targets

The performance of our new method lociPARSE and the other competing methods on 30 independent RNA targets in terms of ground truth as lDDT and RMSD is reported in **Table 1** and **Table 2**, respectively. **Table 1** shows that lociPARSE consistently outperforms all other tested methods across almost all performance criteria when lDDT is used as the ground truth metric. For instance, lociPARSE attains the highest global Pearson’s *r* of 0.67 which is much better than the second-best rsRNAsp (0.5). The same trend continues for global Spearman’s *ρ* (lociPARSE: 0.71 vs. the second-best score of 0.5) and global Kendall’s τ (lociPARSE: 0.55 vs. the second-best score of 0.36. Additionally, lociPARSE attains the lowest diff of 0.06, which is lower than the second-best rsRNASP (0.11). Furthermore, lociPARSE always delivers the highest per-target average correlations. In terms of average lDDT loss, however, DFIRE-RNA attains the lowest average loss (0.05). Meanwhile, lociPARSE, ARES, rsRNASP and RNA3DCNN are tied for the second spot with a comparably low loss of 0.06. To investigate the ability of lociPARSE to distinguish ‘good’ and ‘bad’ models in comparison with the other tested methods across all structural models for all 30 targets, we perform receiver operating characteristics (ROC) analysis using a cutoff of lDDT = 0.75 to differentiate ‘good’ and ‘bad’ models following the CASP15 official assessment [4]. Meanwhile, the area under ROC curve (AUC) quantifies the ability of a method to differentiate ‘good’ and ‘bad’ models. **Table 1** shows that lociPARSE achieves the highest AUC of 0.91, which is noticeably better than second best AUC of 0.8 by ARES, demonstrating better distinguishability aspect of our method.

**Table 1:** Performance on 30 independent RNA targets based on lDDT as the ground truth metric, sorted in nonincreasing order of global Pearson’s *r*. Values in bold indicate the best performance.

| Method | Global |  |  |  |  | Per-target average |  |  |  |
| --- | --- | --- | --- | --- | --- | --- | --- | --- | --- |
| | $r \uparrow$ | $\rho \uparrow$ | $\tau \uparrow$ | Diff ↓ | AUC ↑ | $r \uparrow$ | $\rho \uparrow$ | $\tau \uparrow$ | Loss ↓ |
| lociPARSE | **0.67** | **0.71** | **0.55** | **0.06** | **0.91** | **0.77** | **0.75** | **0.64** | 0.06 |
| rsRNASP | 0.5 | 0.5 | 0.36 | 0.11 | 0.79 | 0.75 | 0.69 | 0.55 | 0.06 |
| RASP | 0.46 | 0.5 | 0.36 | 0.16 | 0.78 | 0.72 | 0.68 | 0.58 | 0.08 |
| ARES | 0.43 | 0.48 | 0.34 | 0.55 | 0.80 | 0.72 | 0.7 | 0.58 | 0.06 |
| DFIRE-RNA | 0.33 | 0.32 | 0.22 | 0.19 | 0.69 | 0.75 | 0.7 | 0.56 | **0.05** |
| cgRNASP | 0.27 | 0.19 | 0.13 | 0.15 | 0.60 | 0.12 | 0.07 | 0.07 | 0.07 |
| RNA3DCNN | 0.19 | 0.17 | 0.12 | 0.64 | 0.58 | 0.47 | 0.36 | 0.27 | 0.06 |

**Table 2:** Performance on 30 independent RNA targets based on all-atom RMSD as the ground truth metric, sorted in nonincreasing order of global Pearson’s *r*. Values in bold indicate the best performance.

| Method | Global |  |  |  | Per-target average |  |  |  |
| --- | --- | --- | --- | --- | --- | --- | --- | --- |
| | $r \uparrow$ | $\rho \uparrow$ | $\tau \uparrow$ | Diff ↓ | $r \uparrow$ | $\rho \uparrow$ | $\tau \uparrow$ | Loss ↓ |
| lociPARSE | **0.65** | **0.65** | 0.47 | 0.31 | **0.59** | **0.61** | **0.49** | **1.47** |
| ARES | 0.64 | 0.64 | **0.48** | 0.3 | 0.58 | **0.61** | 0.48 | 2.0 |
| RASP | 0.64 | **0.65** | 0.47 | **0.21** | **0.59** | 0.56 | 0.44 | 2.37 |
| rsRNASP | 0.63 | **0.65** | 0.47 | 0.26 | 0.52 | 0.52 | 0.37 | 1.93 |
| DFIRE-RNA | 0.54 | 0.53 | 0.39 | 0.22 | 0.49 | 0.49 | 0.35 | 1.88 |
| cgRNASP | 0.49 | 0.48 | 0.33 | 0.29 | 0.18 | 0.09 | 0.09 | 1.59 |
| RNA3DCNN | -0.11 | -0.12 | -0.07 | 0.38 | 0.06 | 0.12 | 0.11 | 2.13 |

Since our method lociPARSE is trained to estimate the lDDT scores, whereas methods such as ARES and RNA3DCNN are trained to estimate the RMSD scores, to ensure a fair performance evaluation, we perform an analogous set of assessment using the all-atom RMSD as the ground truth metric instead of lDDT. As reported in **Table 2**, lociPARSE exhibits remarkable robustness and performance resilience by achieving comparable correlations in both global and per-target levels with the lowest per-target loss, even when evaluated based on RMSD as the ground truth. To further investigate whether each method can effectively sort and rank-order the structures, we analyze the median lDDT and RMSD ground truth metrics of top-1 and best of top-10 ranked structural models from each method. As shown in **Supplementary Figure S1** and **S2**, lociPARSE achieves the highest median top-1 lDDT score of 0.8 and the lowest median top-1 RMSD score of 1.91, demonstrating its effectiveness in identifying the optimal structural model irrespective of the choice of the ground truth metrics. Considering the best of top-10 median scores, lociPARSE is only 0.01 points lower than the highest median lDDT score, and achieves the lowest median RMSD of 1.73. In summary, the results demonstrate that lociPARSE is effective in sorting and rank-ordering structural models while being robust and versatile in terms of the choice of the ground truth assessment metrics.

It is worth mentioning here that both the competing machine learning-based scoring functions ARES and RNA3DCNN exhibit inferior global diff values despite attaining comparable performance in terms of pertarget averages. Meanwhile, DFIRE-RNA, the method attaining the lowest average per-target lDDT loss, does not deliver top performance in terms of global correlations in **Table 1**. That is, there are complementary aspects of scoring and model quality estimation that can lead to performance trade-offs. Our new method lociPARSE strikes an ideal balance to deliver a well-rounded RNA scoring performance across a wide range of assessment metrics. It is interesting to note that among the other tested methods, the two machine learning-based scoring functions ARES and RNA3DCNN show dramatically different performance. While ARES is comparable to the traditional knowledge-based statistical potentials in terms of global correlations and per-target average correlations, RNA3DCNN exhibits poor global and per-target average correlations, which are much lower than most knowledge-based statistical potentials. A similar trend can be observed between rsRNASP and its coarse-grained counterpart cgRNASP, where rsRNASP consistently attains good global and per-target average correlations but cgRNASP falls short. That is, subtle methodological differences such as the granularity of RNA conformational space representation or the choice of the neural network architecture can lead to dramatic differences in performance. Notably, lociPARSE exhibits remarkable robustness by being resilient to the choice of the ground truth assessment metrics or complementary aspects of scoring performance evaluation due to various factors including our novel use of the locality-aware IPA architecture as a general-purpose RNA scoring function, invariant set of features, and the invariant nature of the lDDT metric that lociPARSE is trained to estimate, leading to high-fidelity RNA scoring performance.

### 2.4 Performance on CASP15 RNA targets

**Table 3** and **Table 4** report the performance of lociPARSE and the other competing methods on 12 CASP15 RNA targets based on lDDT and all-atom RMSD as the ground truth metrics, respectively. As shown in **Table 3**, the performance of lociPARSE for the global level assessment based on lDDT as the ground truth metric is considerably better, having the highest global Pearson’s *r* of 0.74, which is much better than the second-best method ARES (0.33). lociPARSE is the second-best method in terms of global diff, only slightly worse by 0.01 points than the lowest global diff. lociPARSE also achieves the highest AUC value of 0.96, outperforming the second-best method rsRNASP (0.83) by a large margin, demonstrating better distinguishability of lociPARSE in separating ‘good’ and ‘bad’ models over a diverse set of predicted structural models submitted by all CASP15 predictors. Furthermore, lociPARSE consistently attains higher per-target average correlations than the other competing methods and achieves the lowest average lDDT loss of 0.07, which is lower than the second-best RNA3DCNN (0.09). **Table 4** reports the full set of results based on all-atom RMSD as ground truth metric that the two competing machine learning-based scoring functions, ARES and RNA3DCNN, are trained on. lociPARSE is better than both methods in terms of both global and per-target correlations with the lowest average RMSD loss, but exhibits comparatively higher global diff. The global correlations of lociPARSE are noticeably better than all competing methods and comparable to most of the energy-based methods in terms of per-target assessment. It is interesting to note that DFIRE-RNA, the method attaining the lowest lDDT loss in 30 independent RNA targets, yields a poor lDDT loss (0.16) in CASP15. By contrast, lociPARSE consistently attains low loss in both test sets, indicating its ability to select the best model that generalizes across different datasets. **Supplementary Figures S1-S2** further demonstrate that the median lDDT score of the top-ranked structure on 12 CASP15 targets by lociPARSE is 0.69, higher than all other methods. In terms of best of top 10 predictions, lociPARSE is second-best with a median lDDT of 0.72 but jointly best in terms of median all-atom RMSD value of 8.29. In summary, the results underscore the ability of lociPARSE to consistently select high-quality structures from a diverse pool of structural models.

**Table 3:** Performance on CASP15 RNA targets based on lDDT as the ground truth metric, sorted in nonincreasing order of global Pearson’s r. Values in bold indicate the best performance.

| Method | Global |  |  |  |  | Per-target average |  |  |  |
| --- | --- | --- | --- | --- | --- | --- | --- | --- | --- |
| | $r \uparrow$ | $\rho \uparrow$ | $\tau \uparrow$ | Diff ↓ | AUC ↑ | $r \uparrow$ | $\rho \uparrow$ | $\tau \uparrow$ | Loss ↓ |
| lociPARSE | **0.74** | **0.72** | **0.55** | 0.12 | textbf0.96 | **0.73** | **0.66** | **0.52** | **0.07** |
| ARES | 0.33 | 0.35 | 0.25 | 0.27 | 0.78 | 0.55 | 0.53 | 0.42 | 0.13 |
| RASP | 0.33 | 0.42 | 0.3 | 0.17 | 0.80 | 0.55 | 0.54 | 0.42 | 0.17 |
| rsRNASP | 0.31 | 0.41 | 0.3 | **0.11** | 0.83 | 0.66 | 0.61 | 0.47 | 0.11 |
| DFIRE-RNA | 0.29 | 0.3 | 0.22 | 0.39 | 0.77 | 0.69 | 0.6 | 0.46 | 0.16 |
| cgRNASP | 0.29 | 0.32 | 0.24 | 0.12 | 0.80 | 0.59 | 0.5 | 0.37 | 0.11 |
| RNA3DCNN | 0 | -0.01 | -0.01 | 0.52 | 0.35 | 0.5 | 0.42 | 0.31 | 0.09 |

**Table 4:** Performance on CASP15 RNA targets based on all-atom RMSD as the ground truth metric, sorted in nonincreasing order of global Pearson’s r. Values in bold indicate the best performance.

| Method | Global |  |  |  | Per-target average |  |  |  |
| --- | --- | --- | --- | --- | --- | --- | --- | --- |
| | $r \uparrow$ | $\rho \uparrow$ | $\tau \uparrow$ | Diff ↓ | $r \uparrow$ | $\rho \uparrow$ | $\tau \uparrow$ | Loss ↓ |
| lociPARSE | **0.37** | **0.34** | **0.24** | 0.58 | 0.33 | 0.32 | 0.24 | **9.74** |
| ARES | 0.15 | 0.11 | 0.09 | 0.23 | 0.22 | 0.25 | 0.18 | 11.83 |
| rsRNASP | 0.13 | 0.21 | 0.14 | 0.49 | **0.34** | **0.37** | **0.27** | 14.37 |
| cgRNASP | 0.11 | 0.13 | 0.09 | 0.41 | 0.28 | 0.31 | 0.23 | 14.79 |
| RASP | 0.1 | 0.09 | 0.07 | 0.63 | 0.18 | 0.2 | 0.15 | 16.79 |
| DFIRE-RNA | 0.06 | 0.11 | 0.09 | 0.86 | 0.32 | **0.37** | **0.27** | 15.31 |
| RNA3DCNN | 0.01 | 0.04 | 0.03 | **0.05** | 0.19 | 0.15 | 0.11 | 13.39 |

When nucleotide-wise quality is evaluated, lociPARSE is orders of magnitude better than RNA3DCNN, the only other method except for lociPARSE that can estimate per-nucleotide score. For example, lociPARSE attains more than three times higher global Pearson’s *r*, Spearman’s *ρ*, and Kendall’s τ than RNA3DCNN, and achieves noticeably lower diff than that attained by RNA3DCNN (**Table 5**). While in principle, the nucleotide-wise scoring performance reported in **Table 5** should be synchronized to the structural level performance reported in **Table 3** as it is the case for our method lociPARSE, RNA3DCNN shows a discrepancy in this regard, possibly due to inconsistencies in nucleotide-wise scoring performance. Overall, lociPARSE delivers stable and consistent nucleotide-wise scoring performance that translates to well-rounded structural level performance.

**Table 5:**
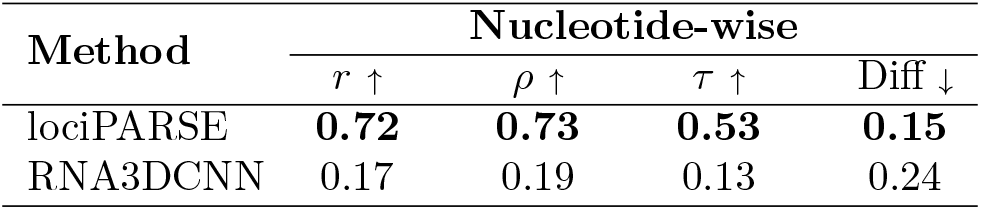
Nucleotide-wise scoring performance on CASP15 set. Values in bold indicate the best performance.

### 2.5 Case study

**Figure 2** shows a representative example of nucleotide-wise scoring performance using lociPARSE for a top-ranked structural model submitted by the winning group AIchemy RNA2 (group 232) for the CASP15 target R1108 having a length of 69. The predicted nucleotide-wise lDDT (pNuL) scores are in close agreement with the ground truth lDDT with a high Pearson’s *r* of 0.89 (**Figure 2a**). Two local problematic regions are estimated by lociPARSE in nucleotide positions (19 − 27) and (59 − 63). These two local problematic regions are visually noticeable when the predicted structural model is aligned with the experimental structure.

**Figure 2:**
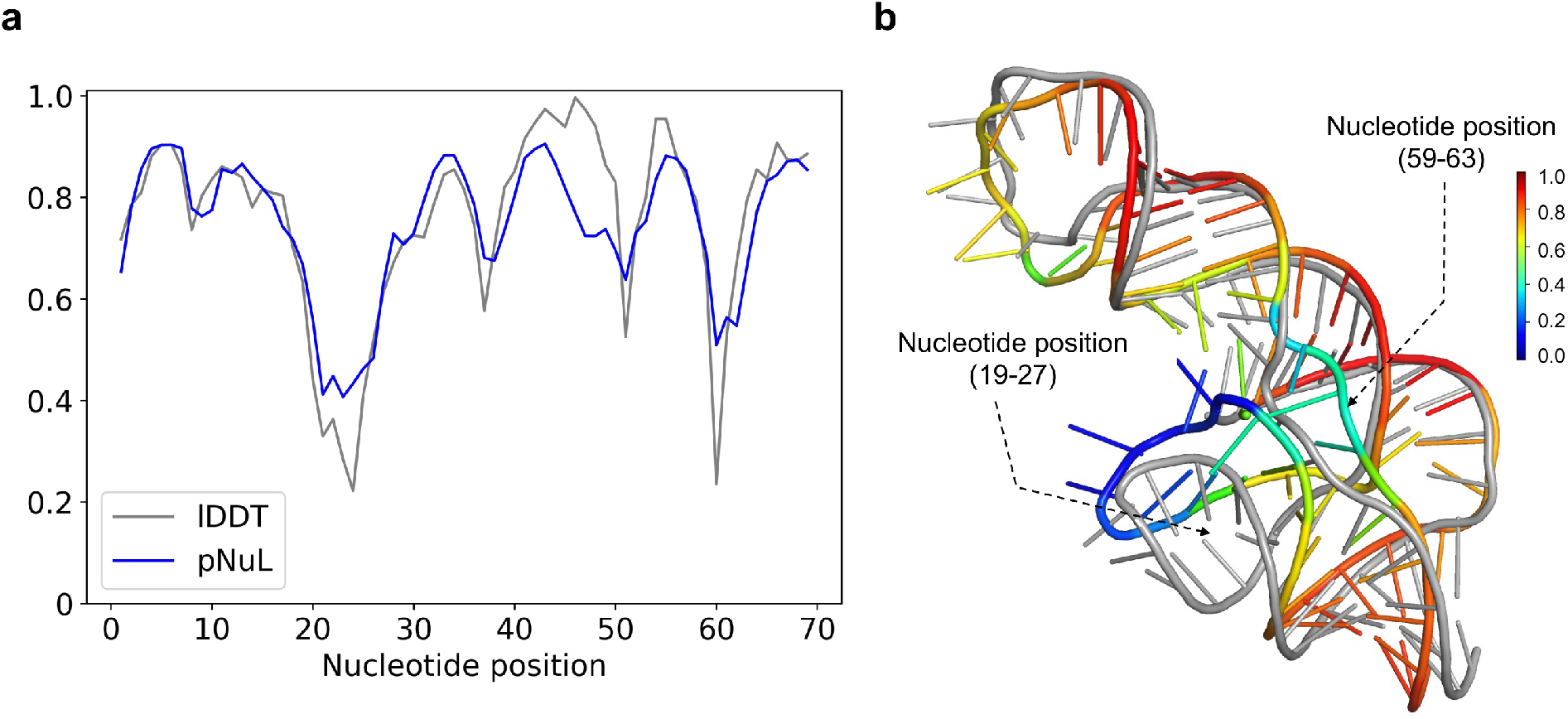
Example of lociPARSE nucleotide-wise quality estimation for the CASP15 target R1108. (a) Predicted nucleotide-wise lDDT (pNuL) vs. the ground truth lDDT for the top-ranked structural model submitted by AIchemy RNA2 (group 232). (b) The predicted structural model in rainbow colored with color code ramping from blue to red for low to high pNuL values superimposed on the experimental structure in gray, and two local problematic regions highlighted.

The poorly modeled structural regions around the hairpin loop in nucleotide positions (19 − 27) and part of the helix strand in positions (59 − 63) are obvious even with simple visual inspection (**Figure 2b**). By contrast, virtually all nucleotides with high pNuL values are structurally well modeled. A comparison with ARES on this target reveals the benefit of nucleotide-wise lDDT prediction as employed by our method over structural level RMSD prediction of ARES, especially in the presence of irregular regions such as the loop in nucleotide positions (19-27). The predicted RMSD value by ARES is 7.8 Å which is noticeably higher than the ground truth all-atom RMSD of 4.63 Å. However, a structural level unfitness score alone fails to pinpoint the problematic regions in the model contributing to the higher RMSD. In contrast, lociPARSE demonstrates its strength by accurately estimating the quality of each nucleotide, effectively identifying the incorrectly modeled loop region in nucleotide positions (19-27) highlighted in blue.

### 2.6 Performance comparison with ARES

To evaluate the effectiveness of our method and perform a more rigorous and head-to-head comparison with ARES free from the influence of the dataset used for training and performance assessment, we retrain our method lociPARSE using the training dataset of ARES and evaluate the performance on the benchmark set 2 from the published work of ARES. Specifically, we retrain lociPARSE using the 16, 000 structural models of 16 RNAs used in ARES (**Supplementary Table S4**) and evaluate the prediction performance of lociPARSE by comparing with ARES on 16 non-redundant RNAs in benchmark set 2 provided by ARES, each containing approximately 5,000 structural models. For performance assessment, we use the all-atom RMSD as the ground truth metric, which ARES is trained to estimate. While the choice of RMSD as the ground truth metric is unfair to our method lociPARSE, which is trained to estimate the lDDT scores, the comparison offers some interesting insights. As shown in **Table 6**, lociPARSE consistently outperforms ARES on almost every aspect, even when using RMSD as the ground truth metric. For example, in terms of both global and per-target assessment, lociPARSE consistently attains better correlations than ARES. While ARES achieves a noticeably lower global diff value compared to lociPARSE likely due to the choice of all-atom RMSD as the ground truth metric that ARES is trained to estimate, lociPARSE outperforms ARES in terms of the per-target average loss, achieving a lower loss value of 8.05. The results highlight the importance of the neural network architecture used in lociPARSE for its improved performance compared to ARES, free from the influence of the dataset used for training and performance assessment, and underscore the robustness and performance resilience of our method.

**Table 6:** Performance comparison on the benchmark set 2 from ARES in terms of all-atom RMSD as ground truth. Values in bold indicate the best performance.

| Method | Global |  |  |  | Per-target average |  |  |  |
| --- | --- | --- | --- | --- | --- | --- | --- | --- |
| | $r \uparrow$ | $\rho \uparrow$ | $\tau \uparrow$ | Diff ↓ | $r \uparrow$ | $\rho \uparrow$ | $\tau \uparrow$ | Loss ↓ |
| lociPARSE | **0.27** | **0.22** | **0.15** | 0.55 | **0.22** | **0.21** | **0.15** | **8.05** |
| ARES | 0.06 | 0.08 | 0.05 | **0.08** | 0 | 0.0 | 0.0 | 9.37 |

### 2.7 Ablation study and hyperparameter selection

To examine the relative importance of the features and architectural hyperparameters adopted in lociPARSE, we conduct ablation experiments by systematically varying individual parameters during model training using the reduced training set and evaluating the accuracy of the independent validation set (Section 4.3). **Table 7** reports the composite quality score (*Q*_*c*_) defined in section 4.5 of the full-fledged version of lociPARSE serving as a baseline and its ablated variants. The results demonstrate that all the parameters adopted in the full-fledged version of lociPARSE positively contribute to the overall accuracy achieved by lociPARSE. For example, we notice a performance decline when we vary the value of K used in the nearest neighbors for the locality computation from K = 20 used in the baseline to K ∈ {5, 10, 30, 40}. Furthermore, to bias the attention weights as well as to update scalar features, we make use of the pair features in the form of nucleotide-nucleotide atomic distances between all pairs of 3 atoms P, C4^*′*^ and N, encoded with Gaussian radial basis functions (hereafter called pair_φ(*d*)_). We notice a significant performance drop when pair_φ(*d*)_ features are isolated as well as changing the number of radial basis functions from RBF = 16 used in the baseline to RBF ∈ *{*1, 2, 4, 8, 32*}*. Similarly, we notice consistent performance decline from the baseline configuration whenever we vary the network architecture such as the number of IPA layers (*N*_*layers*_ = *L*) or various IPA hyperparameters (*N*_*heads*_ = *H, N*_query points_ = *Q, N*_value points_ = V), justifying our choice of the parameters adopted in the full-fledged version of lociPARSE.

**Table 7:** Validation set performance in terms of composite quality score (*Q*_*c*_) with various settings of features (on the left) and hyperparameters (on the right) compared to the full-fledged version of lociPARSE serving as a baseline. Values in bold indicate the best performance.

| Settings | $Q_c$ | Settings | RBF | Hyperparameters | $Q_c$ | Hyperparameters | $Q_c$ |
| --- | --- | --- | --- | --- | --- | --- | --- |
| Baseline<br>(K = 20 w/ pair$_{\varphi(d)}$) | **0.764** | Baseline<br>RBF = 16 | **0.764** | Baseline<br>(L = 4, H = 4) | **0.764** | Baseline<br>(Q = 8, V = 4) | **0.764** |
| K = 5 w/ pair$_{\varphi(d)}$ | 0.744 | RBF = 1 | 0.72 | L = 2 | 0.75 | Q = 4 | 0.762 |
| K = 10 w/ pair$_{\varphi(d)}$ | 0.762 | RBF = 2 | 0.742 | L = 6 | 0.756 | Q = 16 | 0.762 |
| K = 30 w/ pair$_{\varphi(d)}$ | 0.754 | RBF = 4 | 0.74 | H = 2 | 0.748 | V = 2 | 0.746 |
| K = 40 w/ pair$_{\varphi(d)}$ | 0.756 | RBF = 8 | 0.758 | H = 8 | 0.758 | V = 8 | 0.754 |
| K = 20 w/o pair$_{\varphi(d)}$ | 0.728 | RBF = 32 | 0.754 | Q = 2 | 0.754 | V = 16 | 0.758 |

## 3 Discussion

In this work, we developed lociPARSE, a locality-aware invariant point attention model for scoring RNA 3D structures. lociPARSE uses locality information derived from the RNA atomic coordinates to define nucleotide-wise frames together with its local atomic environment. This, coupled with the invariant point attention architecture, allows for the simultaneous estimation of local quality in the form of predicted nucleotide-wise lDDT (pNuL) scores which are then aggregated over all nucleotides by taking an average to estimate global structural correctness in the form of predicted molecular-level lDDT (pMoL). Our empirical results demonstrate the superiority of our method in scoring RNA 3D structures compared to existing approaches.

Our locality-aware attention-based architecture can be extended in several ways, including estimating other local quality measures such as the Interaction Network Fidelity (INF) score [37], which is a local interaction metric that captures various types of base–base interactions in RNA. In fact, INF and lDDT have been shown to correlate well in a near-linear and size-independent relationship [4], suggesting that lDDT may capture the subset of interactions measured in INF whereas INF focuses on a selection of RNA-specific interactions. A model with a very similar architecture as lociPARSE would make an excellent candidate for jointly estimating INF and lDDT, thereby capturing complementary aspects of local quality. Further, a promising direction for future work is to investigate the potential benefits of capturing the multi-state conformational landscape of RNA, since many RNA targets exhibit conformational flexibility [4]. The lDDT score can be computed simultaneously against multiple reference structures of the same RNA at the same time, without arbitrarily selecting one reference structure for the target or removing parts that show variability. Training our model using multi-reference lDDT to capture different classes of conformations will allow our scoring function to account for conformational flexibility and pave the way to evaluate predictions of conformational ensembles instead of just a single structure. One limitation of our method is that it does not account for the stereochemical quality and physical plausibility of the model being evaluated. This is because unlike proteins, the currently available implementation of lDDT for RNA does not penalize for stereochemical violations. Using a customized version of lDDT that incorporates stereochemical quality checks in its calculation can address such limitations, and this aspect remains an important future direction.

## 4 Methods

### 4.1 Model input

Our model uses only input features derived directly from a nucleotide sequence and RNA 3D structural coordinates. We use just the basic nucleotide-level encodings for our input. These include one-hot encoding of the nucleotide (i.e., a binary vector of 5 entries indicating each of the 4 nucleotide types and one for non-standard nucleotides such as ‘T’ or modified nucleotides) and the relative position of the nucleotide in its sequence calculated as *i*/*L* (where i is the nucleotide index and *L* is the sequence length). For our pair features, we use the index i of a nucleotide’s partner in sequence and the corresponding 3D coordinates, quantified by sequential separation and spatial proximity information. We do not consider the nucleotide type of the partner as a pair feature. The sequence separation i.e., the absolute difference between the two nucleotide indices is discretized into 5 bins and represented by one-hot encoding where the first two bins correspond to self-loops and adjacent bonds respectively. The rest of the three bins are defined based on three types of interactions depending on the sequence separation: short-range (2-5), medium-range (6-24) and long-range (>24), similar to [38, 39]. The other component of our pair features includes nucleotide-nucleotide atomic distances between all pairs of P, C4^*′*^ and glycosidic N atoms, encoded with radial basis functions. The radial basis functions are defined based on the distance from a reference point, making them suitable for capturing distance-based similarities. We used Gaussian radial basis functions to encode the inter-atomic distances in the following manner. For a nucleotide pair (*i, j*), at first *k* = 9 distances 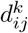 are calculated because of all possible combinations among the set of 3 atoms P, C4^*′*^ and N. Then, the set of all 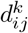 values, *D* is encoded using Gaussian radial basis function as follows:

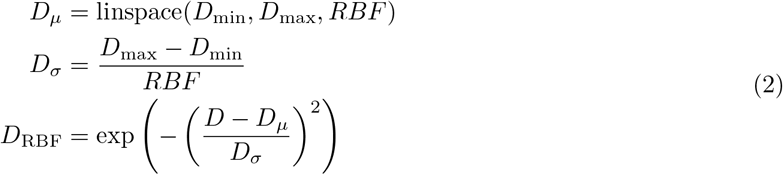

where, RBF = 16 is the number of radial basis functions, a value chosen based on the ablation experiment presented in **Table 7** on an independent validation set. As such, *D*_*µ*_ is a set of 16 linearly spaced values between *D*_min_ = 0 and *D*_max_ = 100, which indicates the minimum and maximum inter-atomic distances, respectively. The output *D*_RBF_ contains 16 encoded distance pair features for each distance 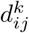. It is important to note that all our nucleotide and pair features are invariant, consistent with the invariant layers of the IPA module.

### 4.2 Network architecture

#### 4.2.1 Construction of local nucleotide frames

To perform invariant point attention on a set of 3D points, we represent each nucleotide in a geometric abstraction using the concept of frames. Each nucleotide frame in the form of a tuple is defined as a Euclidean transform 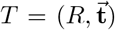, where *R* ∈ *ℝ*^3*×*3^ is a rotation matrix and 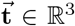 is the translation vector that can be applied to transform a position in local coordinates 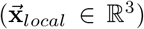 to a position in global coordinates 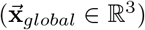 as:

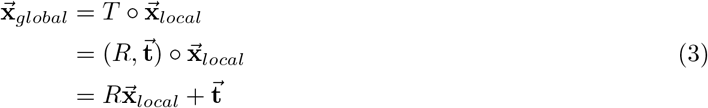

In our setting, we define local nucleotide frames from the Cartesian coordinates of P, C4^*′*^, and glycosidic N atoms of the input RNA 3D structure and construct 3-bead coordinate frame using a Gram–Schmidt process specified in Alphafold2 (Algorithm 21) that takes the input coordinates (scaled by 0.1) of P as 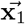, C4^*′*^ as 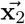, and N as 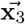. Note that the translation vector 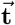 is assigned to the center atom 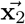.

#### 4.2.2 Locality-aware invariant point attention

The formulation of locality-aware IPA used in our work combines sequence representation, ***s***_*i*_, from each nucleotide i of the input RNA, pair representation ***e***_*ij*_ of nucleotide *i* with other nucleotides *j* based on nucleotide pair adjacencies capturing the local atomic environment 𝒩 of nucleotide *i*, where *j* ∈ 𝒩_*i*_ is the locality information derived from the RNA atomic coordinates. Consequently, the update function of the IPA layer is as follows:

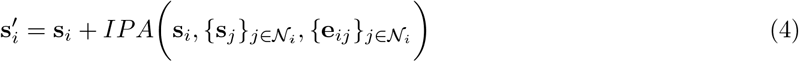

To perform attention on 3D point clouds, IPA derives query 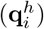, key 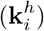 and value 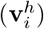 embeddings from a linear projection of **s**_*i*_ to a latent representation of dimension c for each nucleotide i, where 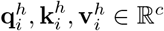 and *h* ∈ *{*1, …, *N*_*head*_*}* which represents the number of attention heads in the IPA module. 3D query, key and value points are also generated considering the local frame T_i_ of each nucleotide i, where 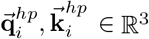, p ∈ {1, …, *N*_query points_*}* and 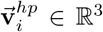, *p* ∈ {1, …, *N*_query points_}. The IPA module acts on a set of frames (parameterized as Euclidean transforms of the local frame T_*i*_) and is invariant under global Euclidean transformations *T*_*global*_ on said frames. By performing locality-aware geometry and edge-biased attention, the IPA module transforms the 3D points from the target nucleotide’s local frame into a global reference frame for computing the attention weights as follows:

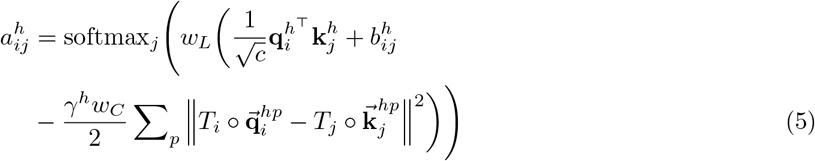

where 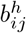 is the attention bias derived from the linear projection of **e**_*ij*_ to hidden dimension *c*, weighting factors *w*_*L*_ and *w*_*C*_ are taken from the IPA formulation specified in AlphaFold2 and *γ*^*h*^ ∈ ℝ is a learned scalar value. The attention mechanism acting on a set of local frames ensures invariance under global Euclidean transformations such as global rotations and translations of the input RNA due to the invariant nature of *ℓ*_2_-norm of a vector under such rigid transformations.

The attention weights are used to compute the outputs of the attention mechanism while mapping them back to the local frame and preserving invariance, as follows:

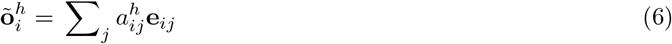

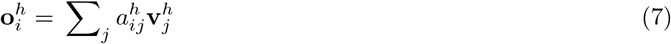

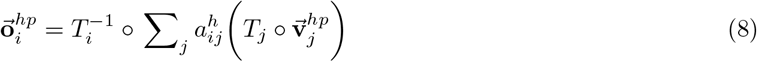

The outputs of the attention mechanism are then concatenated and passed through a linear layer to compute the updated sequence representations 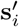 of each nucleotide as follows:

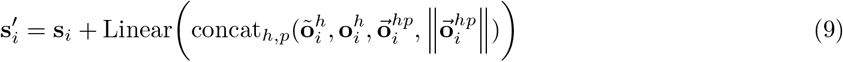

The updated sequence embeddings 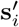 for each nucleotide i are subsequently stacked together to obtain the embedding ***s***^*′*^ for all nucleotides in the RNA. Finally, a linear layer followed by a 2 layer fully connected network implemented as a multilayer perceptron (MLP) is used to obtain the final representation ***s***^*f*^ before estimating nucleotide-wise lDDT scores as follows:

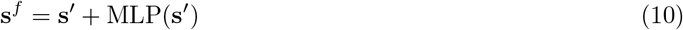

### 4.3 Training, validation and test datasets

To curate our training dataset, we first obtained the training dataset used in trRosettaRNA [12] containing 3, 632 RNA targets. We then filtered this set by removing duplicate chains and discontinuous structures, separating monomers from complexes, splitting multiple chains into single chains, and correcting formatting issues in the coordinates files. We removed sequences with length > 200 nucleotides and ensured that our training and test sets were non-redundant by running CD-HIT-est [40] with default parameter settings, which reduced the training set to 1, 399 RNA targets. We generated 37 structural models per target with a total of 51, 763 structural models for the 1, 399 targets using a combination of different RNA 3D structure prediction tools including recent deep learning-enabled RNA structure prediction methods [11–15], physics-based RNA folding [34], and structure perturbation using PyRosetta [35]. For the deep learning-enabled RNA structure prediction methods, we used default parameter settings to generate the structural models. Additionally, we generated one model using SimRNA, selecting the first frame of the cluster with a 3.5 Å RMSD threshold. Finally, each of the 6 structures generated by DeepFoldRNA [11] was relaxed by FastRelax protocol [41] within the PyRosetta framework [35] with 5,000 and 10,000 steps aiming to introduce structural diversity in our training set. The number of structural models generated by each method is listed in **Supplementary Table S1**. In addition, we separately curated a validation set of 60 RNA targets for the ablation study and hyperparameter selection from the Protein Data Bank (PDB) [42] with experimental structures released between January 1, 2022 and July 6, 2023. Such a date range was chosen to avoid any overlap with our training dataset collected from trRosettaRNA which used structures released before January 1, 2022. Details of the curation process for this validation set is available in [42]. We generated 25 structural models for each of these 60 RNAs as well as for the 30 independent RNAs in the first test set Test30 using the recent deep learning-based RNA structure prediction methods [11–15]. Supplementary Tables S2 and **S3** list the number of structural models per target used for our two test sets Test30 and CASP15, respectively. The nucleotide-wise ground truth lDDT distributions of our training and test sets are shown in **Supplementary Figures S3-S4**. We created a reduced training subset consisting of 6, 872 structural models for 1, 399 RNA targets through clustering [43] for ablation study and hyperparameter selection.

### 4.4 Training details

To train our model, lociPARSE, we obtained nucleotide-wise ground truth lDDT scores by comparing the predicted structural models in our training dataset against the corresponding experimental structures using the docker version of OpenStructure [44] available here. During the ground truth lDDT computation, we enabled the option ‘–lddt-no-stereochecks’ to skip stereochemical quality checks in its calculation following the recent CASP assessment in [4]. lociPARSE was implemented in PyTorch [45] with the ℒ1 loss function to learn the mean absolute error between ground truth lDDT and predictions on nucleotide level, thereby formulating the local nucleotide-wise quality estimation as a regression task. We trained our model using the Adam optimizer [46] having parameters *β*1 = 0.9 and *β*2 = 0.999 with a learning rate of 0.0001 and dropout rate of 0.1. The training process consists of 50 epochs on an 80-GB NVIDIA A100 GPU.

### 4.5 Competing methods and evaluation metrics

lociPARSE is compared against both traditional knowledge-based statistical potentials (rsRNASP [20], RASP [22], DFIRE-RNA [23], and cgRNASP [21]) and recent machine learning-based RNA scoring functions (RNA3DCNN [17] and ARES [31]). rsRNASP is an all-atom distance-dependent potential considering short and long-ranged interactions present in RNA based on sequence separation aiming to capture the hierarchical nature of RNA folding [47]. Ribonucleic Acids Statistical Potential (RASP) is another all-atom statistical potential based on the averaging reference state. Similar to rsRNASP, RASP also separates interaction pairs into local and non-local categories and takes into account the base stacking and base pairing interactions present in RNA. DFIRE-RNA is yet another distance-scaled statistical potential designed using a finite-ideal-gas reference state. Finally cgRNASP, a coarse-grained counterpart of rsRNASP potential introduces three different variants of coarse-grained potentials for RNA scoring. We have used the 3-bead representation of cgRNASP in this work which takes into account P, C4^*′*^ and N atoms. The Atomic Rotationally Equivariant Scorer (ARES) is an equivariant graph neural network that scores RNA structures by identifying complex structural motifs through equivariant convolutions. ARES employs E3NN [48] to predict the global RMSD of the structure. Finally, RNA3DCNN uses 3D convolutional neural network to predict the RMSD-like unfitness score of a nucleotide to its surroundings by considering RNA 3D structure as a 3D image and representing each nucleotide as an array of voxels. For prediction, we used the model that was trained on samples generated from both molecular dynamics (MD) and Monte Carlo (MC) simulations. It is worth noting that except lociPARSE, RNA3DCNN is the only other method that estimates both local and global quality.

Our assessment metrics include both global and average per-target Pearson’s (r), Spearman’s rank (ρ), and Kendall’s Tau rank (τ) correlation coefficients between the estimated score of each method and the ground truth lDDT as well as ground truth all-atom RMSD where a higher correlation indicates better performance. While Pearson’s r assumes a linear relationship between variables and is sensitive to outliers, Spearman’s *ρ* and Kendall’s τ account for non-linear but monotonic trends and are less affected by outliers. Also, Kendall’s τ is particularly robust for small sample sizes and many tied ranks. Therefore, reporting all three coefficients provides a comprehensive assessment of the predicted quality scores of different methods. Ground truth lDDT of the test set targets are calculated using the docker version of OpenStructure [44] whereas the ground truth all-atom RMSD is calculated using casp-rna pipeline [4]. As discussed in section 2.2, all the predicted quality scores and the ground truth metrics are normalized between (0-1] during scoring performance evaluation. Diff, another assessment metric is calculated at the global level as the mean absolute difference between the structural level estimated scores (pMoL) of all the structures of all targets and their corresponding ground truth lDDT or RMSD metrics. Per-target loss or top-1 loss is calculated as the absolute difference between the ground truth lDDT or RMSD of the structural model ranked at the top by each method and the ground truth lDDT or RMSD of the most accurate structural model for each target which is then averaged over all targets to get the loss value for each method over a test set. Lower values of diff and loss, therefore, indicate better performance. To calculate the local nucleotide-wise scoring performance in **Table 5**, we accumulated all the nucleotide-wise predicted scores (pNuL) and computed the coefficients and diff in the same manner as in our global assessment. For RNA3DCNN, we adhered to the method described in their work to scale the nucleotide-level predicted RMSD in the range (0-1] before the evaluation. We additionally performed receiver operating characteristics (ROC) analysis using a lDDT threshold of 0.75 to separate ‘good’ and ‘bad’ structural models, following [4]. Consequently, the area under the ROC curve (AUC) quantifies the ability of a scoring function to distinguish good and bad models. Finally, an average of all assessment metrics is taken to combine the results of all the different metrics into a single composite quality score, called *Q*_*c*_, defined as:

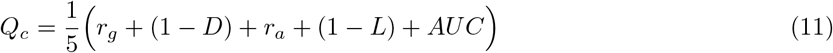

where, *r*_*g*_ = global Pearson’s *r, D* = global diff, *r*_*a*_ = per-target average Pearson’s r, L = average loss, and AUC = area under the ROC curve. We use the composite quality score for ablation study and hyperparameter selection, where higher values of *Q*_*c*_ indicate better performance.

## 5 Acknowledgements

This work was partially supported by the National Institute of General Medical Sciences (R35GM138146 to D.B.) and the National Science Foundation (DBI2208679 to D.B.).

