## Supplementary Information for "lociPARSE: a locality-aware invariant point attention model for scoring RNA 3D structures"

### 1 Supplementary Tables

#### 1.1 Training dataset

We generated 37 decoys for each of the 1,399 RNA targets using various deep learning techniques, physics-based RNA folding methods, and PyRosetta perturbation methods. The total number of structures in our training set is 51,730. The training set was divided into two subsets: a training set and a validation set, with a split ratio of 80:20. The training set was used to train the architecture, while the validation set was used to save the model based on the best loss value which is used for evaluation throughout. The RNA 3D structural models and their corresponding ground truth lDDTs used to train and validate lociPARSE with a detailed description can be found here [1].

Table S1: List of the number of decoys per RNA target in our in-house training set

| Method | Number of decoys |
|---|---|
| DeepFoldRNA | 6 |
| trRosettaRNA | 10 |
| RoseTTAFoldNA | 1 |
| DRfold | 6 |
| RhoFold | 1 |
| SimRNA | 1 |
| DeepFoldRNA + PyRosetta | 12 |
| Total | 37 |

#### 1.2 Test datasets

##### 1.2.1 Test30 and hyperparameter optimization set

We generated 25 decoys for each target in Test30 and hyperparameter optimization dataset TS60. So the total number of structures in the Test30 set is 750. The decoy structures of CASP15 were collected from the CASP15 website which consists of all the submitted models in CASP15 by all groups. All the structural models for the three datasets can be found here [1].

Table S2: Number of decoys per RNA target in our in-house benchmark sets.

| Method | Number of decoys per target |
|---|---|
| DeepFoldRNA | 6 |
| trRosettaRNA | 10 |
| RoseTTAFoldNA | 1 |
| DRfold | 7 |
| RhoFold | 1 |
| Total | 25 |

##### 1.2.2 CASP15 test set

Table S3: List of the number of decoys per RNA target in CASP15 test set

| Target | Number of decoys per target |
|---|---|
| R1107 | 131 |
| R1108 | 115 |
| R1116 | 145 |
| R1117 | 153 |
| R1126 | 140 |
| R1128 | 137 |
| R1136 | 158 |
| R1138 | 130 |
| R1149 | 138 |
| R1156 | 145 |
| R1189 | 136 |
| R1190 | 132 |
| Total | 1660 |

#### 1.3 ARES training set

We excluded structural models of 255d and 283d from the set since we couldn't find the experimentally determined native structure for these two targets. The four targets have been used as validation set following [2] to save the trained model based on best validation loss: 1q9a, 1kka, 1i9x and 1a4d.

Table S4: List of targets from ARES's training set to train lociPARSE

| Structure IDs | |
|---|---|
| 157d | 1mhk |
| 1a4d | 1q9a |
| 1csl | 1qwa |
| 1dqf | 1xjr |
| 1esy | 28sp |
| 1i9x | 2a43 |
| 1kd5 | 2f88 |
| 1kka | 1l2x |

### 2 Supplementary Figures

#### 2.1 Ability to rank the structures in terms of lDDT and RMSD

2.1.1 and 2.1.2 shows the performance of all the quality assessment methods in terms of the ability to sort the pool of structures for each target in our 2 benchmark test sets Test30 and CASP15 following the analysis of top deep learning RNA quality assessment method in literature ARES [2].

##### 2.1.1 Top 1 median score

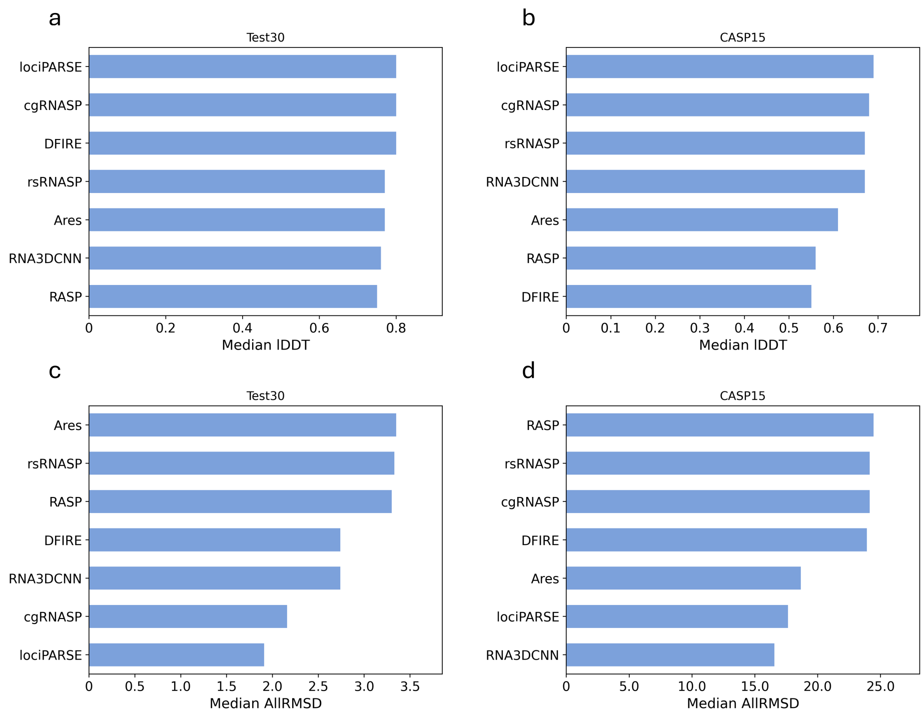

Figure S1: (a) and (b) represents the median lDDT for targets in Test30 and CASP15 datasets respectively in terms of the top ranked structure for each target assessed by 7 different methods. (c) and (d) shows the corresponding median all atom RMSD for same analysis.

##### 2.1.2 Top 10 median score

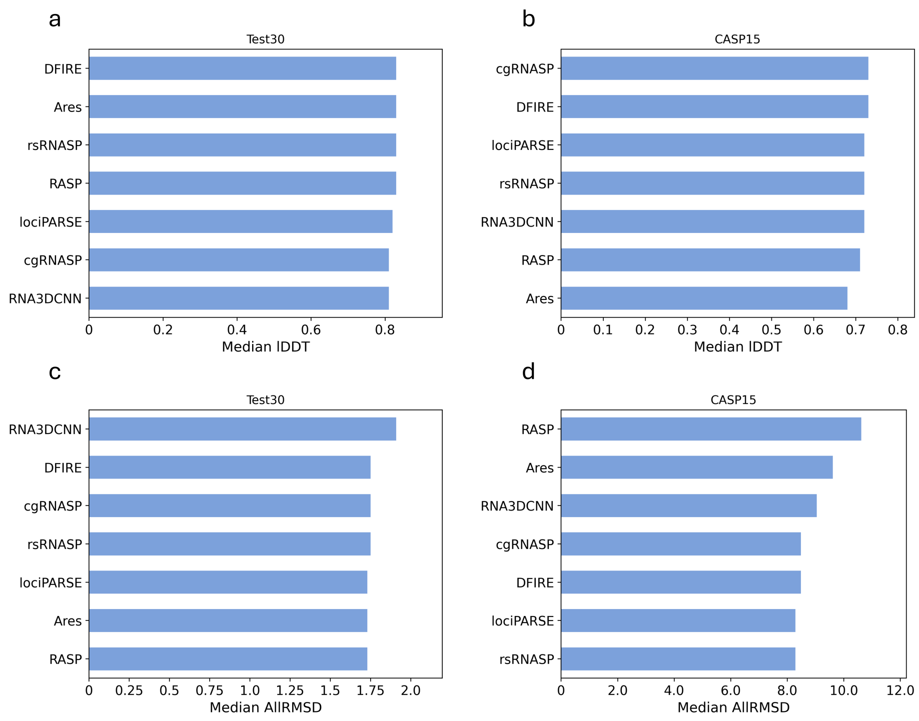

Figure S2: (a) and (b) represents the median lDDT for targets in Test30 and CASP15 datasets respectively in terms of the highest lDDT among 10 best scoring structural models for each target assessed by 7 different methods. (c) and (d) shows the corresponding all atom RMSD for same analysis.

#### 2.2 Training dataset distributions

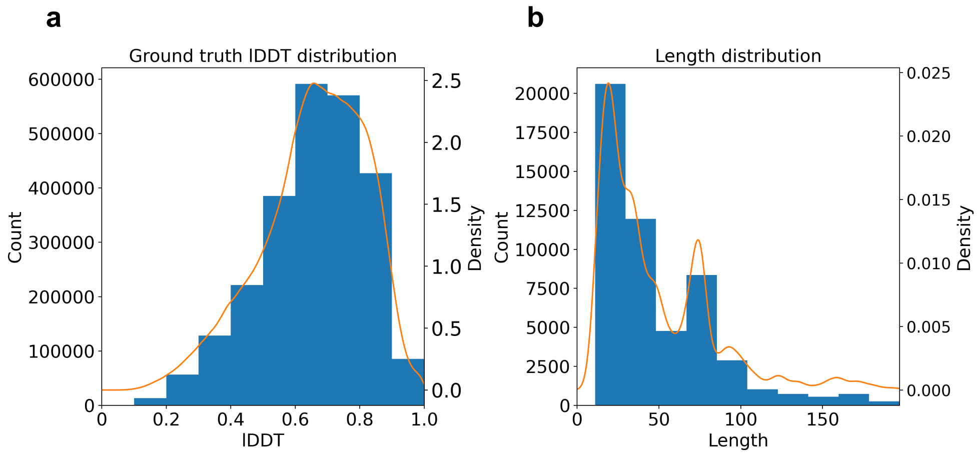

Figure S3: (a) Nucleotide-wise ground truth lDDT distribution of approximately 2.5 million nucleotides on our training set. (b) Length distribution of our training set.

#### 2.3 Test dataset distributions

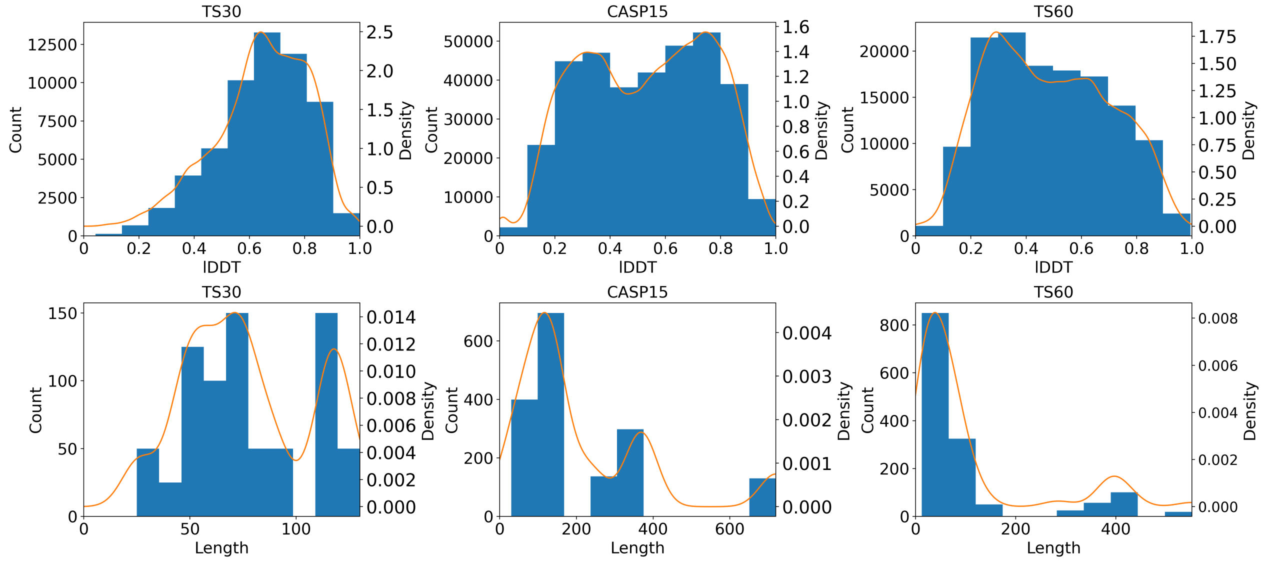

Figure S4: Nucleotide-wise ground truth lDDT and corresponding length distributions for the test and validation sets.
